# Genetically increased serum calcium levels reduce Alzheimer’s disease risk

**DOI:** 10.1101/255059

**Authors:** Qinghua Jiang, Yang Hu, Shuilin Jin, Guiyou Liu

**Affiliations:** School of Life Science and Technology, Harbin Institute of Technology, Harbin, China; Department of Mathematics, Harbin Institute of Technology, Harbin, China

**Keywords:** Alzheimer’s disease, serum calcium, genome-wide association study, Mendelian randomization

## Abstract

**IMPORTANCE** Alzheimer’s disease (AD) is the leading cause of disability in the elderly. It has been a long time about the calcium hypothesis of AD on the basis of emerging evidence since 1994. However, most studies focused on the association between calcium homeostasis and AD, and concerned the intracellular calcium concentration. Only few studies reported reduced serum calcium levels in AD. Until now, it remains unclear whether serum calcium levels are genetically associated with AD risk.

**OBJECTIVE** To evaluate the genetic association between increased serum calcium levels and AD risk

**DESIGN, SETTING, AND PARTICIPANTS** We performed a Mendelian randomization study to investigate the association of increased serum calcium with AD risk using the genetic variants from the large-scale serum calcium genome-wide association study (GWAS) dataset (N=61,079 individuals of European descent) and the large-scale AD GWAS dataset (N=54,162 individuals including 17,008 AD cases and 37,154 controls of European descent). Inverse-variance weighted meta-analysis (IVW) was used to provide a combined estimate of the genetic association. Meanwhile, we selected the weighted median regression and MR-Egger regression as the complementary analysis methods to examine the robustness of the IVW estimate.

**EXPOSURES** Genetic predisposition to increased serum calcium levels

**MAIN OUTCOMES AND MEASURES** The risk of AD.

**RESULTS** We selected 6 independent genetic variants influencing serum calcium levels as the instrumental variables. IVW analysis showed that a genetically increased serum calcium level (per 1 standard deviation (SD) increase 0.5-mg/dL) was significantly associated with a reduced AD risk (OR=0.56, 95% CI: 0.34-0.94, P=5.00E-03). Meanwhile, both the weighted median estimate (OR=0.60, 95% CI: 0.34-1.06, P=0.08) and MR-Egger estimate (OR=0.66, 95% CI: 0.26-1.67, P=0.381) were consistent with the IVW estimate in terms of direction and magnitude.

**CONCLUSIONS AND RELEVANCE** We provided evidence that genetically increased serum calcium levels could reduce the risk of AD. Meanwhile, randomized controlled study should be further conducted to assess the effect of serum calcium levels on AD risk, and further clarify whether diet calcium intake or calcium supplement, or both could reduce the risk of AD.

**Key Points:** **Question** Is there a genetic relationship between elevated serum calcium levels and the risk of Alzheimer’s disease?

**Findings** This Mendelian randomization study showed that the genetically increased serum calcium levels were associated with the reduced risk of Alzheimer’s disease.

**Meaning** These findings provide evidence that genetically increased serum calcium levels could reduce the risk of Alzheimer’s disease.

## Introduction

Calcium is involved in many biological processes including the many neural processes in the body ^1-2^. Altered neuronal calcium homeostasis is widely hypothesized to underlie cognitive deficits in normal aging subjects, and many neurodegenerative diseases including Alzheimer’s disease (AD) ^2-6^. A polymorphism rs2986017 in CALHM1 influences calcium ion homeostasis, beta amyloid levels, and AD risk ^7^.

Until now, most studies focused on the association between calcium homeostasis and AD, and concerned the intracellular calcium concentration ^2,4-7^. The involvement of intracellular calcium dysregulation in AD has been well established ^2^. A recent article updated the calcium hypothesis of AD on the basis of emerging evidence since 1994 ^8^. This updated hypothesis provided a framework for integrating new evidence such as aging, genetic, and environmental factors into a comprehensive theory of pathogenesis ^8^. To develop more effective therapies, we should establish the causal links between the clinical and neurobiological phenotypes of dementia and neurodegeneration ^8^. Mendelian randomization method using large-scale genome-wide association study (GWAS) datasets could avoid some limitations of observational studies, and is widely used to determine the causal inferences ^1,9-11^. However, there is no GWAS to identify the causal variants influencing the calcium homeostasis or intracellular calcium levels. Until now, it is still impossible to detect the genetic relationships of calcium homeostasis or intracellular calcium levels with AD risk.

In addition to the calcium homeostasis and intracellular calcium levels in AD, evidence also highlighted the role of serum calcium levels in AD risk. Landfield et al. found that AD cases were characterized by lower serum calcium levels compared with age-matched controls or demented subjects with mild indications of vascular contributions ^12^. The serum chemistry tests further showed that moderately low values of phosphate, calcium, or both identified 74% of AD cases and 100% of early onset AD cases, compared with only 46% of mixed/vascular dementia cases and 31% of normal age-matched control samples ^12^. Compared with mildly affected individuals, the severely demented patients had lower serum calcium levels ^13^. Meanwhile, compared with normal age-matched controls, AD patients showed a decrease in serum calcium level, increase in serum parathyroid hormone level, increase in urinary calcium excretion, and a decrease in serum 1, 25 dihydroxyvitamin D (1,25(OH)2D) concentration ^14^. However, it remains unclear whether serum calcium levels are genetically associated with AD risk.

In recent years, large-scale GWAS promptly identified some common genetic variants and provided insight into the genetics of serum calcium and AD ^15-16^. The existing large-scale GWAS datasets provide strong support for investigating the potential genetic association between serum calcium levels and AD risk by a Mendelian randomization analysis method. Here, we hypothesize that there may be a genetic relation between serum calcium levels and AD risk. We performed a Mendelian randomization study to investigate the potential genetic association using a large-scale serum calcium GWAS dataset and a large-scale AD GWAS dataset.

## Materials and methods

### Study Design

Mendelian randomization is based on the premise that the human genetic variants are randomly distributed in the population ^17^. These genetic variants are largely not associated with confounders and can be used as instrumental variables to estimate the causal association of an exposure (serum calcium levels) with an outcome (AD) ^17^. The Mendelian randomization is based on three principal assumptions (Figure 1), which have been widely described in recent studies ^1,17^. First, the genetic variants selected to be instrumental variables should be associated with the exposure (serum calcium levels) (assumption 1 in Figure 1) ^1,17^. Second, the genetic variants should not be associated with confounders (assumption 2 in Figure 1) ^1,17^. Third, genetic variants should affect the risk of the outcome (AD) only through the exposure (serum calcium levels) (assumption 3 in Figure 1) ^1,17^. The second and third assumptions are collectively known as independence from pleiotropy ^17^. This study is based on the publicly available, large-scale GWAS summary datasets. All participants gave informed consent in all these corresponding original studies.

**Figure 1.**
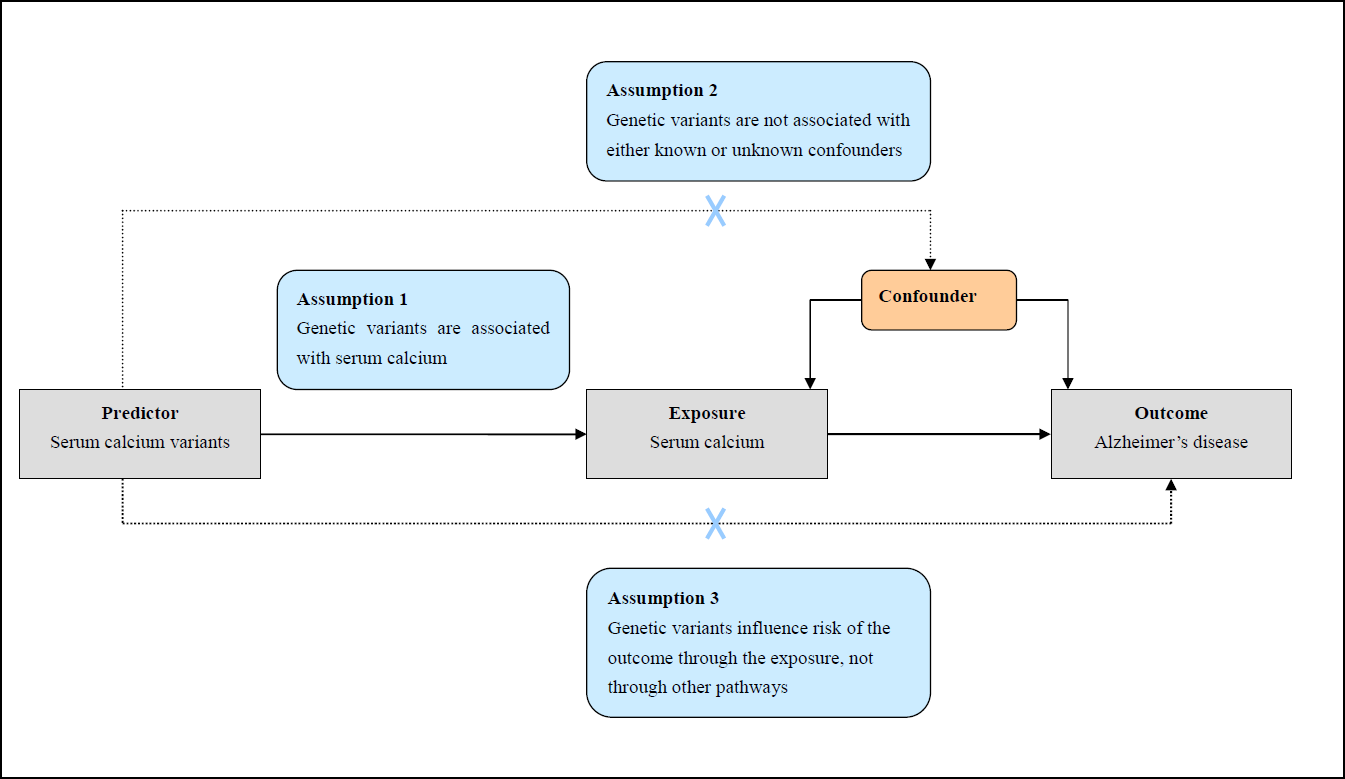
Mendelian randomization assumptions The Mendelian randomization is based on three principal assumptions, which have been widely described in recent studies ^1,17^. First, the genetic variants selected to be instrumental variables should be associated with the exposure (serum calcium levels) (assumption 1) ^1,17^. Second, the genetic variants should not be associated with confounders (assumption 2) ^1,17^. Third, genetic variants should affect the risk of the outcome (AD) only through the exposure (serum calcium levels) (assumption 3) ^1,17^.

### Serum Calcium GWAS Dataset

Here, we selected genetic variants that influence serum calcium levels as the instrumental variables based on the GWAS dataset of serum calcium concentration ^15^. This GWAS included 39400 individuals from 17 population-based cohorts in discovery stage and 21679 individuals in replication stage (N=61079 individuals of European descent) ^15^. The discovery stage and the meta-analysis of the discovery and replication stage identified 8 genetic variants to be associated with serum calcium levels with the genome-wide significance (*P* < 5.00E-08) ^15^. All these 8 genetic variants were located in different genes and were not in linkage disequilibrium (Table 1) ^15^. We provided more detailed information about the methods to measure serum calcium levels in the eTable 1.

### AD GWAS Dataset

The AD GWAS dataset is from the large-scale meta-analysis, which is performed by the International Genomics of Alzheimer’s Project (IGAP) ^16^. In stage 1, the IGAP genotyped and imputed 7,055,881 SNPs, and performed a meta-analysis of four GWAS datasets including 17,008 cases and 37,154 controls of European descent ^16^. All patients with AD satisfied the NINCDS-ADRDA criteria or DSM-IV guidelines ^16^. Here, we extracted the summary statistics of these 8 genetic variants with AD risk in stage 1 dataset.

### Pleiotropy Analysis

In Mendelian randomization study, one important issue is potential violation of assumption 2 and 3 through pleiotropy occurring when a genetic instrument is associated with a study outcome through biological pathways outside the exposure of interest. Here, we performed an assessment for pleiotropy to assure that the selected genetic variants do not exert effects on AD risk through biological pathways independent of serum calcium levels. A number of steps were taken to reduce the risk of pleiotropy. Here, we provided more detailed information about five stages of pleiotropy analysis in eMethods.

### Mendelian Randomization Analysis

Until now, three different Mendelian randomization analysis methods have been well established including inverse-variance weighted meta-analysis (IVW), weighted median regression and MR-Egger regression ^1,9-11,17-20^. Here, we selected the IVW as the main analysis method to perform the Mendelian randomization analysis, as did in previous studies ^1,9-11,17-20^. Meanwhile, we selected the weighted median regression and MR-Egger regression as the complementary analysis methods to examine the robustness of the IVW estimate. The weighted median regression method could derive consistent estimates when up to 50% of instruments are not valid ^17^. MR-Egger could provide a statistical test the presence of potential pleiotropy, and account for this potential pleiotropy ^19-20^. If there is no clear evidence of pleiotropy, we would expect all three tests to give consistent estimates. The odds ratio (OR) as well as 95% confidence interval (CI) of AD corresponds to per 0.5-mg/dL increase (about 1 standard deviation (SD)) in serum calcium levels. All analyses were conducted using the R package ‘MendelianRandomization’ ^21^. The threshold of statistical significance for the potential genetic association between serum calcium levels and AD risk was *P* < 0.05. We also performed a sensitivity analysis using the leave-one-out permutation method, and a power analysis. Here, we provided more detailed information about the IVW, weighted median, MR-Egger regression, sensitivity analysis and power analysis in the eMethods.

## Results

### Genetic variant validation

In addition to statistical association, all these 8 genetic variants could map to genes implicated in calcium pathways or related phenotypic traits ^15^. In brief, GCKR, DGKD, CASR, GATA3, DGKH/KIAA0564, CYP24A1, and CARS were associated with bone metabolism and endocrine control of calcium ^15^. More detailed information has been described in previous study ^15^. The physiological function of VKORC1L1, a paralogous enzyme sharing about 50% protein identity with VKORC1, is unknown. However previous findings showed that VKORC1 could inhibit calcium oxalate salt crystallization, adhesion and aggregation ^22^.

### Association of serum calcium variants with AD

Of these 8 genetic variants associated with serum calcium levels, we successfully extracted the summary statistics for 7 variants in stage 1 dataset of AD GWAS. The rs17711722 is not available in stage 1 dataset of AD GWAS. We then selected its proxy rs1829942, which showed high LD with rs17711722 (r²=0.91 and D’=0.96) using the HaploReg v4.1 based on LD information in 1000 Genomes Project (CEU) with r^2^ >= 0.8^23^. None of these 8 genetic variants was significantly associated with AD risk at the Bonferroni corrected significance threshold (*P*<0.05/8=0.0063) (Table 1).

**Table 1.**
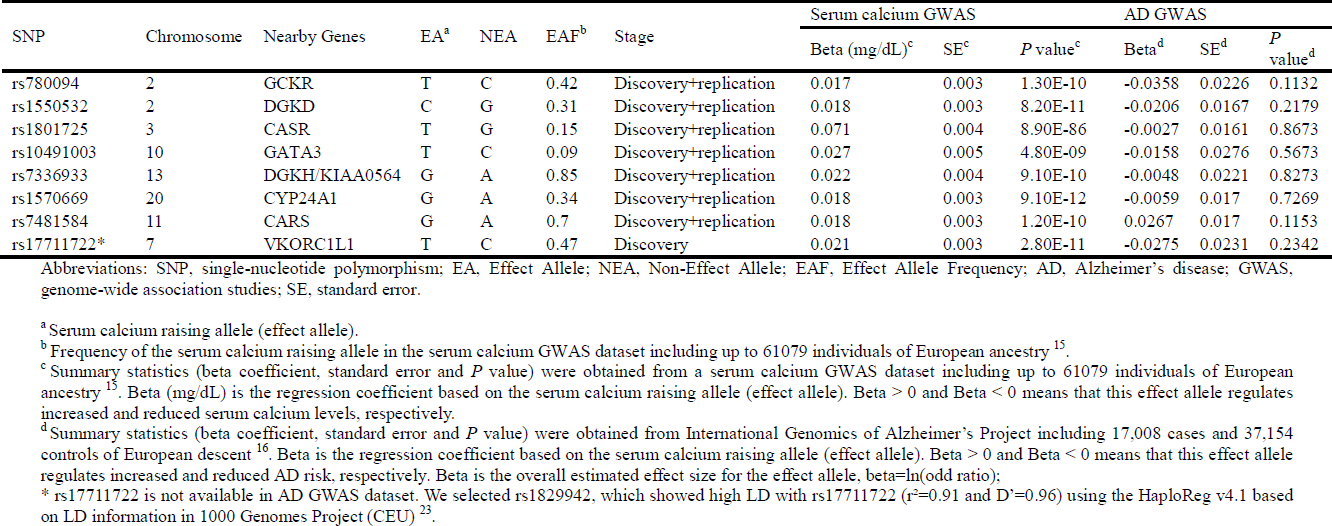
Characteristics of 8 genetic variants in serum calcium and AD GWAS datasets

### Pleiotropy analysis

In stage 1, rs780094 was significantly associated with known confounders at the Bonferroni corrected significance threshold (*P*<0.05/8=0.0063), as described in eTable 2. In brief, rs780094 variant was significantly associated with type II diabetes (*P*=1.00E-05), hip circumference (*P*=3.40E-05), and waist hip ratio adjusted for BMI (*P*=1.80E-03). In stage 2, we found that none of these 8 variants was significantly associated with dietary factors including mineral supplements and vitamin supplements. However, two genetic variants were significantly associated with alcohol drinking or alcohol dependence at the Bonferroni corrected significance threshold (*P*<0.05/8=0.0063), including rs780094 with alcohol drinking (*P*=3.65E-09), and rs7481584 with alcohol dependence (*P*=9.50E-05 and alcohol dependence early age of onset (*P*=4.70E-04). More detailed information is provided in eTable 3.

In stage 3, none of these 8 genetic variants showed significant association with Parkinson’s disease ^24^, AD biomarkers ^25^, and AD age at onset survival ^26^ at the Bonferroni corrected significance threshold (*P*<0.05/8=0.0063) (eTable 4). In stage 4, we found evidence that rs780094 had pleiotropic associations with lipids, glycemic traits, type 2 diabetes, and measures of adiposity at the Bonferroni corrected significance threshold, which have been identified in a recent study ^1^. No variant was significantly associated with circulating 25-OH vitamin D concentrations (all *P*>0.05) in the SUNLIGHT consortium ^15^.

In stage 5, both heterogeneity test and MR-Egger intercept test showed no significant heterogeneity (I^2^=0%, 95% CI [0%; 59.1%], and *P=*0.5937) and significant intercept (MR-Egger intercept β=0.007; *P*=0.527) in these 8 genetic variants, respectively. However I^2^ had a wider 95% CI [0%; 59.1%] including the maximum I^2^=59.1%, which means that the potential heterogeneity has not been fully excluded. By iteratively pruning the corresponding variant list, we found that there was no heterogeneity in the remaining 7 genetic variants by excluding rs7481584 with I^2^=0% and 95% CI [0%; 0%] (Table 2).

**Table 2.**
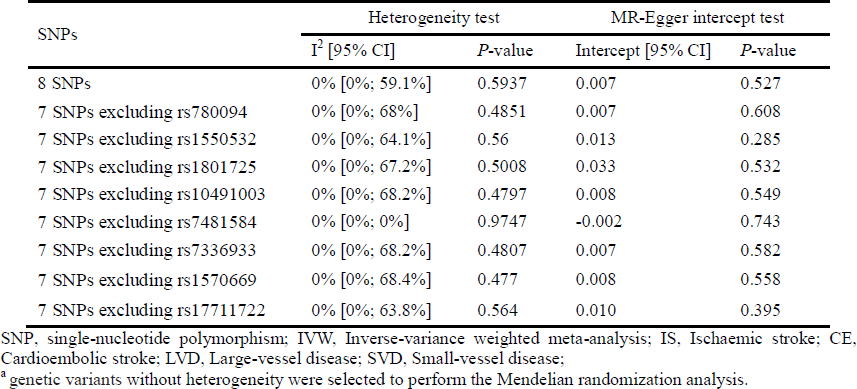
pleiotropy analysis using heterogeneity test and MR-Egger intercept test

In summary, our pleiotropy analysis showed that rs780094 and rs7481584 may have potential violation of assumption 2 and 3 through pleiotropy. To meet the Mendelian randomization assumptions (Figure 1), we excluded both variants in following analysis.

### Association of serum calcium with AD risk

Using the remaining 6 genetic variants, IVW analysis showed that a genetically increased serum calcium level (per 1 SD increase 0.5-mg/dL) was significantly associated with a reduced AD risk (OR=0.56, 95% CI: 0.34-0.94, *P*=5.00E-03) (Table 3). Interestingly, both the weighted median estimate (OR=0.60, 95% CI: 0.34-1.06, *P*=0.08) and MR-Egger estimate (OR=0.66, 95% CI: 0.26-1.67, *P*=0.381) were consistent with the IVW estimate in terms of direction and magnitude. Meanwhile, both heterogeneity test and MR-Egger intercept test showed no significant heterogeneity (I^2^=0%, 95% CI [0%; 0%], and *P=*0.957) and significant intercept (MR-Egger intercept β=-0.006; *P*=0.403) between these 6 variants. Figure 2 shows individual genetic estimates from each of the 6 genetic variants using different methods.

**Figure 2.**
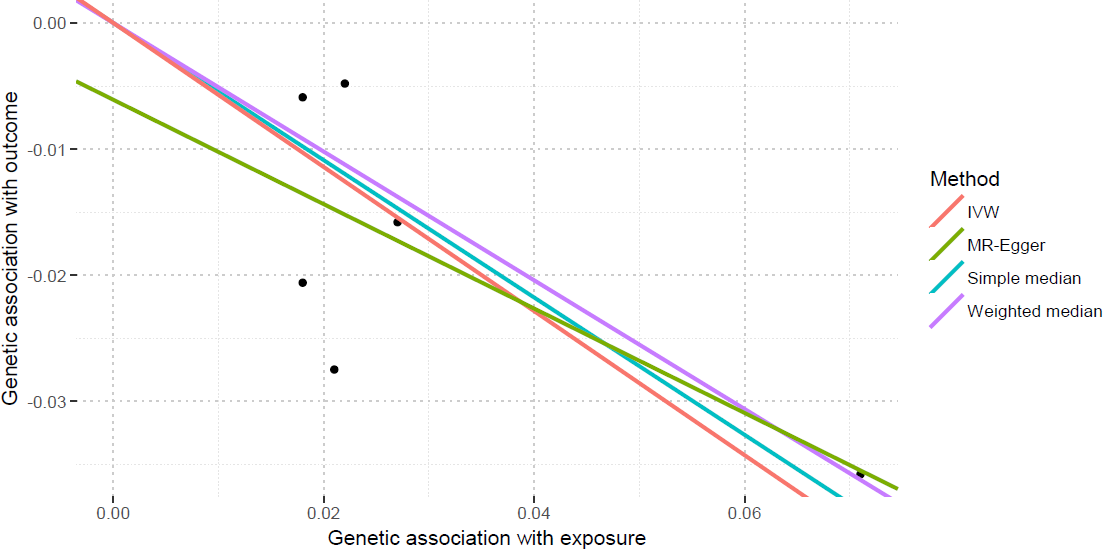
Individual genetic estimates from each of 6 genetic variants using different methods This scatter plot show individual causal estimates from each of 6 genetic variants associated with serum calcium levels on the x-axis and AD risk on the y-axis. The continuous line represents the causal estimate of serum calcium levels on AD risk.

To further test the stability of these estimates, we sequentially removed each genetic variant in the Mendelian randomization analysis. The direction and precision of the genetic estimates between increased serum calcium levels and reduced risk of AD remained largely unchanged using all these three Mendelian randomization methods (Table 3).

**Table 3.**
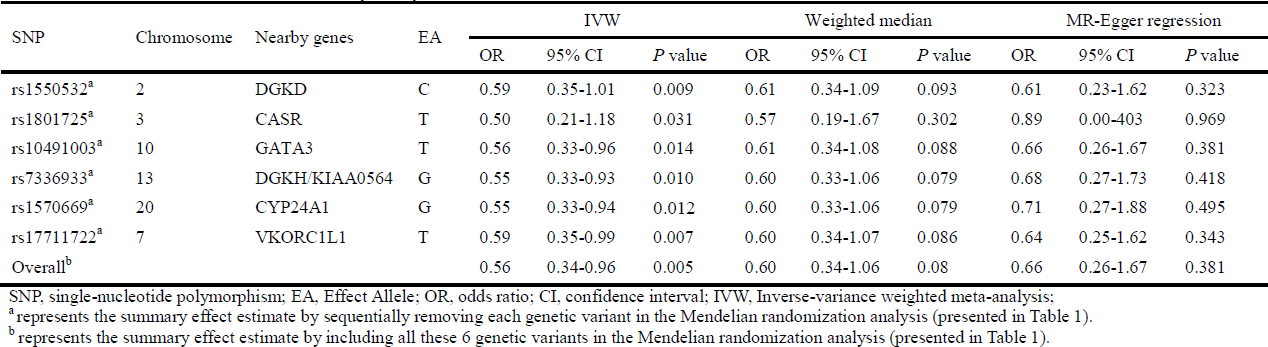
sensitivity analysis of the association between increased serum calcium levels and AD risk

### Power analysis

We further performed a power analysis using the strength of the instrumental variables (6 genetic variants) R^2^ 0.81%, the actual N for AD GWAS 54,162, and the proportion of cases 0.314. Interestingly, the power is 100% to detect the genetic association between increased serum calcium levels and reduced AD risk with OR=0.56. The N required for 80% power is 16427.

## Discussion

Until now, it has been a long time about the calcium hypothesis of AD on the basis of emerging evidence since 1994 ^8^. However, most studies focused on the association between calcium homeostasis and AD, and concerned the intracellular calcium concentration ^2,4-7^. Only few studies reported reduced serum calcium levels in AD ^12-14^. Until now, it remains unclear whether serum calcium level (diet and supplements) is genetically associated with AD risk. Fortunately, the existing large-scale serum calcium and AD GWAS datasets prompts us to investigate the potential genetic association between serum calcium and AD risk by a Mendelian randomization method. Our findings indicated genetically increased serum calcium levels were significantly associated with a reduced AD risk. Each SD higher serum calcium level (1SD=0.5 mg/dL) was associated with a 44% reduced AD risk (OR=0.56, 95% CI: 0.34-0.94, *P*=5.00E-03).

Interestingly, our results are consistent with previous studies, which reported reduced serum calcium levels in AD ^12-14^. Meanwhile, a recent study also reported the reduced serum calcium levels in Parkinson’s disease ^27^. The levels of serum calcium in Parkinson’s disease group with dementia were significantly lower than the normal control group (*P*<0.001) ^27^. There is also a correlation between serum calcium levels and cognitive impairment ^27^. Lower serum calcium levels could predict worse cognitive scores ^27^.

More than 99% of calcium in human body is stored in the skeleton as hydroxyapatite contributing to skeletal strength ^28^. Only less than 1% of calcium is located in the blood, soft tissues, and extracellular fluid ^28^. About 51% serum calcium exists in the free or ionized state ^28^. It is the ionized calcium that could enter cells and cause the activation of essential physiological processes ^28^. Calcium homeostasis is a complex process involving multiple key components. Normal calcium homeostasis could be regulated by three major hormones including parathyroid hormone release governed by the calcium-sensing receptor, calcitonin, and the hormonally active form of vitamin D (1,25(OH)2D), which act on their corresponding receptors in gut, kidney, and bone, respectively ^15^. Meanwhile, calcium homeostasis could also be regulated by ionized serum calcium ^28^. Importantly, two recent Mendelian randomization studies provided clear evidence that increased vitamin D could reduce the risk of AD and another neurodegenerative disease multiple sclerosis ^10,29^. Hence, our results are comparable to these recent findings.

The 2016 AD facts and figures showed an estimated 5.4 million people to have AD in United States ^30^. By 2050, the number of people living with AD will be 13.8 million in the United States ^30^. Until now, two ways could increase the serum calcium levels. One is diet calcium intake. Until now, the recommended daily calcium intake is 1000 to 1200 mg ^31^. It is difficult to get this recommended amount through diet alone, so calcium supplements are widely used ^31^. In the United States, about 43% of people, including about 70% of older women, take calcium supplements ^32^. Hence, the genetic association between serum calcium levels and AD risk may have clinical and public health implications.

Until now, our findings should be considered to be exploratory, as did in observational studies. In order to translate these genetic findings into clinical and public health implications, the potential mechanisms underlying this genetic association remain to be thoroughly evaluated. Meanwhile, randomized controlled study should be conducted to assess the effect of serum calcium levels on AD risk, and further clarify whether diet calcium intake or calcium supplement, or both could reduce the risk of AD.

### Strengths

This Mendelian randomization study has several strengths. First, this study may benefit from the large-scale serum calcium GWAS dataset (N=61079 individuals) and AD GWAS dataset (N=54,162 individuals), which provide ample power (100%) to detect the genetic association between serum calcium levels and AD risk. Second, both the serum calcium and AD GWAS datasets are from the European descent, which may reduce the influence on the potential association caused by the population stratification. Third, multiple independent genetic variants are taken as instruments, which may reduce the influence on the potential association caused by the linkage disequilibrium. Fourth, we selected multiple Mendelian randomization methods, which increase the precision of the estimate. Fifth, we performed a pleiotropy analysis to exclude two genetic variants associated with potential confounders, which meets the Mendelian randomization assumptions.

### Limitations

First, we only selected one AD GWAS dataset. We think that a replication data may be necessary. However a replication dataset with a similar large-scale of AD GWAS dataset was not available. Second, five steps were taken to reduce the risk of pleiotropy, as did in our pleiotropy analysis. However, we could not completely rule out additional confounders. Until now, it is almost impossible to fully rule out pleiotropy present in any Mendelian randomization study ^1,17,33^. Third, it could not be completely ruled out that population stratification may have had some influence on the estimate. In order to reduce the effect of population stratification, our study was restricted to individuals of European ancestry. Fourth, the genetic association between serum calcium levels and AD risk may differ by ethnicity or genetic ancestry. This genetic association should be further evaluated in other ancestries.

Fifth, Mendelian randomization can be extended to conduct a mediation analysis, estimating the proportion of an observed association of an exposure with an outcome that occurs through a given mediator ^17^. Until now, there is no GWAS to evaluate the calcium channel blockers. Hence, we could not conduct the medication analysis, as the medication data is not available now. In our further work, we will further conduct the medication analysis, when the medication data is publicly available.

Sixth, polygenic risk score (PRS) is another measure of genetic predisposition ^17^. PRS is conceptually similar to Mendelian randomization ^34^. There are also some differences ^34^. Mendelian randomization study just uses two summary GWAS datasets. The PRS method uses one summary GWAS dataset, and one genotype data. Until now, a large-scale AD GWAS dataset with the individual genotype data is not publicly available. In our further work, we will further conduct a PRS analysis, when the individual genotype data is publicly available.

## Conclusions

In summary, we demonstrate that genetically increased serum calcium levels could reduce AD risk in people of European descent using large-scale GWAS datasets. These findings provide rationale for further investigating the clinical and public health implications of high serum calcium levels by diet or supplement, or both in preventing the progression of AD.

## Acknowledgments

This work was supported by the Major State Research Development Program of China (No: 2016YFC1202302), and the National Natural Science Foundation of China (No: 61571152).

